# LizardNet: A mobile hybrid deep learning tool for classification of 3D representations of Amazonian lizards

**DOI:** 10.1101/2024.01.15.575627

**Authors:** Arthur Gonsales da Silva, Roger Pinho de Oliveira, Caio de Oliveira Bastos, Elena Almeida de Carvalho, Bruno Duarte Gomes

## Abstract

Image classification is a highly significant field in machine learning (ML), especially when applied to address longstanding and challenging issues in the biological sciences. In this study, we present the development of a hybrid deep learning-based tool suitable for deployment on mobile devices. This tool is aimed at processing and classifying three-dimensional samples of endemic lizard species from the Amazon rainforest. The dataset used in our experiment was collected at the Museu Paraense Emílio Goeldi (MPEG), Belém-PA, Brazil, and comprises three species: a) *Anolis fuscoauratus*; b) *Hoplocercus spinosus*; and c) *Polychrus marmoratus*. We compared the effectiveness of four artificial neural networks (ANN) for feature extraction: a) MobileNet; b) MobileNetV2; c) MobileNetV3Small; and d) MobileNetV3Large. Additionally, we evaluated five classical ML models for classifying the extracted patterns: a) Support Vector Machine (SVM); b) GaussianNB (GNB); c) AdaBoost (ADB); d) K-Nearest Neighbors (KNN); and e) Random Forest (RF). Our most effective model, MobileNetV3-Small + Linear SVM, achieved an accuracy of 0.948 and a f1-score of 0.955. Notably, it not only proved to be the least complex model among all combinations but also demonstrated the best performance after a statistical comparison. These results indicate that the combination of deep learning (DL) models with less complex classical ML algorithms, which have a lower error propensity, emerges as a viable and efficient technique for classifying three-dimensional lizard species samples. Such an approach facilitates taxonomic identification work for professionals in the field and provides a tool adaptable for integration into mobile data recording equipment, such as smartphones.

**Author summary:** The taxonomic classification of lizards requires an exceptional level of knowledge and attention to minute details beyond the ordinary to accurately categorize specimens. Such tasks impose significant mental and visual costs on humans, unlike computer vision algorithms capable of extracting visual patterns from images imperceptible to the human eye. In this research, we utilized a dataset from the herpetarium of the Emílio Goeldi Museum in Belém-PA, Brazil. The data were self-captured, with each sample comprised of three photos: dorsal, lateral, and ventral views of each specimen. The sample size was constrained by the quality and abundance of preserved specimens, necessitating the application of a data augmentation method on the pre-separated training and validation sets. This augmentation led to a considerable increase in the number of samples per species, from a few dozen to several hundred. Our experimental approach involved utilizing pre-trained neural networks to extract 3D sample characteristics, subsequently classified using classical machine learning algorithms. This hybrid strategy was adopted due to the nature of data collection and synthetic data augmentation. Our method enables specimen identification through three-dimensional representations, allowing for a more comprehensive utilization of morphological information by the model.

## Introduction

In the Squamata order, which comprises species that, among other characteristics, have their bodies covered by scales, the classification of lizards is based on multiple morphological features [1]. According to [2], these morphological characteristics are referred to as microornamentations and are most prominent in the dorsal scales of the head, trunk, and tails of each individual. Modern biodiversity data collection equipment, such as sound recorders, camera traps, and other imaging methods, allow the measurements of many parameters that make possible the extraction of vast amounts of information in a relatively inexpensive manner. This technology has become increasingly popular among scientists and helps to answer questions such as: a) Which species occur in a given area?; b) What are their activities/behaviour?; and c) How many individuals inhabit the region? [3]. The success in inventorying and monitoring forest lizard species relies on robust monitoring and sampling and currently represents one of the most complex tasks in the field of herpetological conservation [4].

One of the most used data types in problems involving biodiversity conservation with specialized image models is camera trap images [5]. The aim of remote monitoring can range from species identification to inferring the abundance and distribution of important conservation animals, but these motivations typically share a common goal - to classify target species [6]. This interest in remote monitoring is accompanied by several challenges in large-scale identification [6].

The most recent research in automated identification of animal species can be divided into two distinct types: laboratory-based investigation (LBI), and field-based investigation (FBI) [7]. For LBI, a pre-established image acquisition protocol must be followed to standardize the sampling and use of specimens, which are typically handled by a specialized biologist. This contrasts significantly with FBI, where a mobile device or camera is usually employed for the image acquisition process of the individuals [7].

In studies of insect classification, for instance, LBI is the most commonly used method due to the highly manual handling of specimens [40]. On the other hand, the identification of mammals and fish is typically accomplished using field-recorded images, while automated recognition of plant species can benefit from both the controlled environment of a laboratory and field conditions [8]. These studies focus on the use of Machine Learning (ML) with Convolutional Neural Networks (CNN), which are models specialized in image processing that extracts high-level abstractions from data and are considered the state-of-the-art for tasks involving image classification [9].

The most common type of algorithm learning used for image classification is supervised learning, where input data (samples) are fed into the model along with their corresponding labels (class names), and the algorithms are trained to map the input information to the output label, such as the name of a species, for example [16].

Before the emergence of computer vision (CV) models and artificial intelligence (AI) algorithms in general, the process of identifying and conserving animal species was and still is, in some places, carried out manually with a high dependence on human activities, which imposes several limitations on the task [15]. These limitations, mainly physical and cognitive, hinder the understanding of species distribution and diversity. For instance, counting of colonies of seabirds and cave-dwelling bats conducted by humans tends to significantly underestimate the actual number of individuals [15]. This scenario of limitations and uncertainties changed with the advent of large-scale AI-driven automation of these tasks.

With recent advances in automated image classification and information gathering, new approaches have become possible [40]. Several existing examples demonstrate the applications of automatic classification based on deep learning (DL) using taxonomic data from different species [9]. Table 1 summarizes recent studies where CV algorithms were employed to perform automated species identification [8, 10–12, 15].

**Table 1.** Recent studies where computer vision algorithms were employed for species classification in different taxonomic groups.

|  | Samples | Architecture | Accuracy | Study |
| --- | --- | --- | --- | --- |
| *Reptiles* | 386,006 | Vision Transformer (ViT) | 0.962 | Bhardwaj, Manish, et al. (2023) |
| *Reptiles* | 82,601 | EfficientNet | 0.870 | Durso, Andrew M., et al. (2021) |
| *Reptiles & Amphibians* | 2,700 | VGG16 | 0.870 | Binta Islam, Sazida, et al. (2023) |
| *Fishes* | 1,080 | Image Processing + SVM | 0.942 | Sharmin, Israt, et al. (2019) |
| *Mammals* | 326 | Mask R-CNN + ResNet101 | 0.980 | Gray, Patrick C., et al. (2019) |

As can be seen in table 1, most studies used pre-trained models. This is the case because when pre-trained networks are employed either as feature extractors or efficiently optimized for the new dataset, there exists a strong correlation between the high accuracy achieved by the model on its original pre-training phases with its score in the new training demand [14]. Thus, incremental or transfer learning only requires the pre-trained model to generalize an additional predictive pattern that might be present in the dataset while retaining its previous optimal weights often gathered on ImageNet Large-Scale Visual Recognition Competition (ILSVRC) [11].

In this study, we have developed an open-source system for the automatic classification of three-dimensional samples of Amazonian lizard species, adapted for deployment on mobile equipment such as smartphones. We employed state-of-the-art DL and ML techniques for image processing and classification using the family of CNNs known as MobileNets [26–28], together with classical ML models, which demonstrated exceptional efficiency in similar tasks. Despite the widespread use of CNNs in taxonomic databases [8, 10–12, 15], our reviews revealed no applications of these models, or hybrids of these models, to three-dimensional specimens of Amazonian lizards. We validated our model using synthetic data generated from the previously separated training and test sets, as well as original images from the collection at Museu Paraense Emílio Goeldi (MPEG).

## Results

### Dataset complexity & model performance

We processed one copy of the image dataset with each variant of the MobileNet network, and it proved to be a crucial strategy in determining the optimal classifier. The complexity of each dataset played a fundamental role in the performance of classical ML algorithms. Figure 1 below illustrates the difference in the clustering for each dataset as revealed by t-distributed Stochastic Neighbor Embedding (t-SNE) [33].

**Fig 1.**
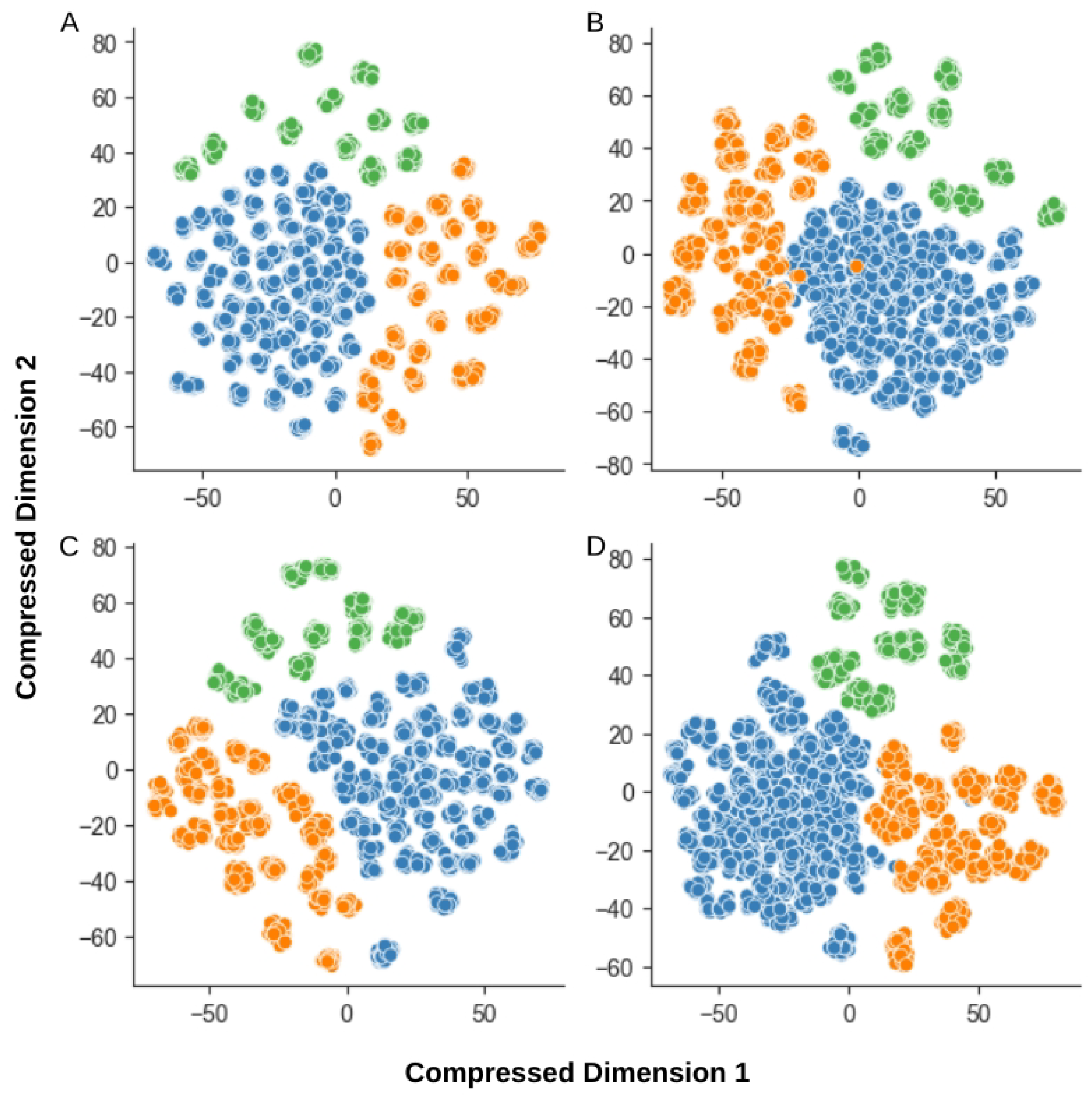
The t-SNE analysis of each full-features dataset. (a) MobileNet (b) MobileNetV2 (c) MobileNetV3-Large (d) MobileNetV3-Small.

Analyses (a) and (c) exhibit good spacing between clusters, but the samples are more dispersed among themselves. Analysis (b) shows a more apparent class overlap, despite each cluster being relatively well concentrated. Analysis (d), obtained from the data extracted with MobileNetV3-Small, presents the best trade-off between cluster separation and sample concentration, with little to no apparent class overlap. Based on the analysis using t-SNE, as expected, the impact of the complexity of each dataset is determinant for the model performance. Table 2 presents the top-performing models trained with all features extracted by the variants of the MobileNet.

**Table 2.** Classic ML models performances on each full-features dataset generated by each MobileNet variant.

|  | Best Model | Average F1-Score | Average Accuracy |
| --- | --- | --- | --- |
| MobileNetV3-Small | Linear SVM | 0.973 | 0.974 |
| MobileNetV2 | Linear SVM | 0.970 | 0.963 |
| MobileNet (V1) | Linear SVM | 0.961 | 0.970 |
| MobileNetV3-Large | Linear SVM | 0.942 | 0.953 |

The combination of MobileNetV3-Small + Linear SVM produced a model that outperformed the others trained with all features. Table 3 shows the same comparison for the models trained with the 20 top-ranked features only.

**Table 3.**
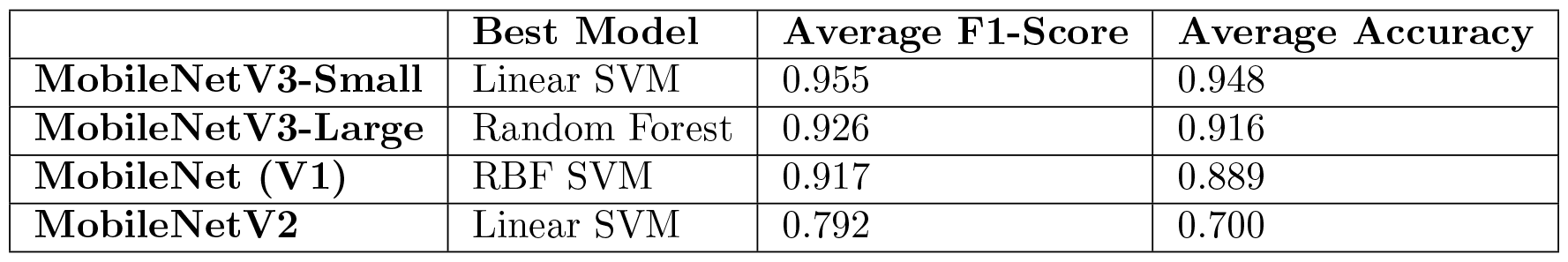
Classic ML models performances on each 20 top-ranked features dataset generated by each MobileNet variant.

### Models’ performance statistical evaluation

The McNemar’s statistical test, which compares the confusion matrix of two algorithms with paired samples [39] was conducted on the MobileNetV3-Small + Linear SVM models for both full-features and 20 top-ranked features datasets, and resulted in a Chi-squared value of 9.0 and a p-value of 1.0, which suggests that both models have statistically the same performance. This ensures the safe utilization of the least complex one. The Figure 2 shows the confusion matrix of the best model trained with the 20 top-ranked features, evaluated on it’s validation set.

**Fig 2.**
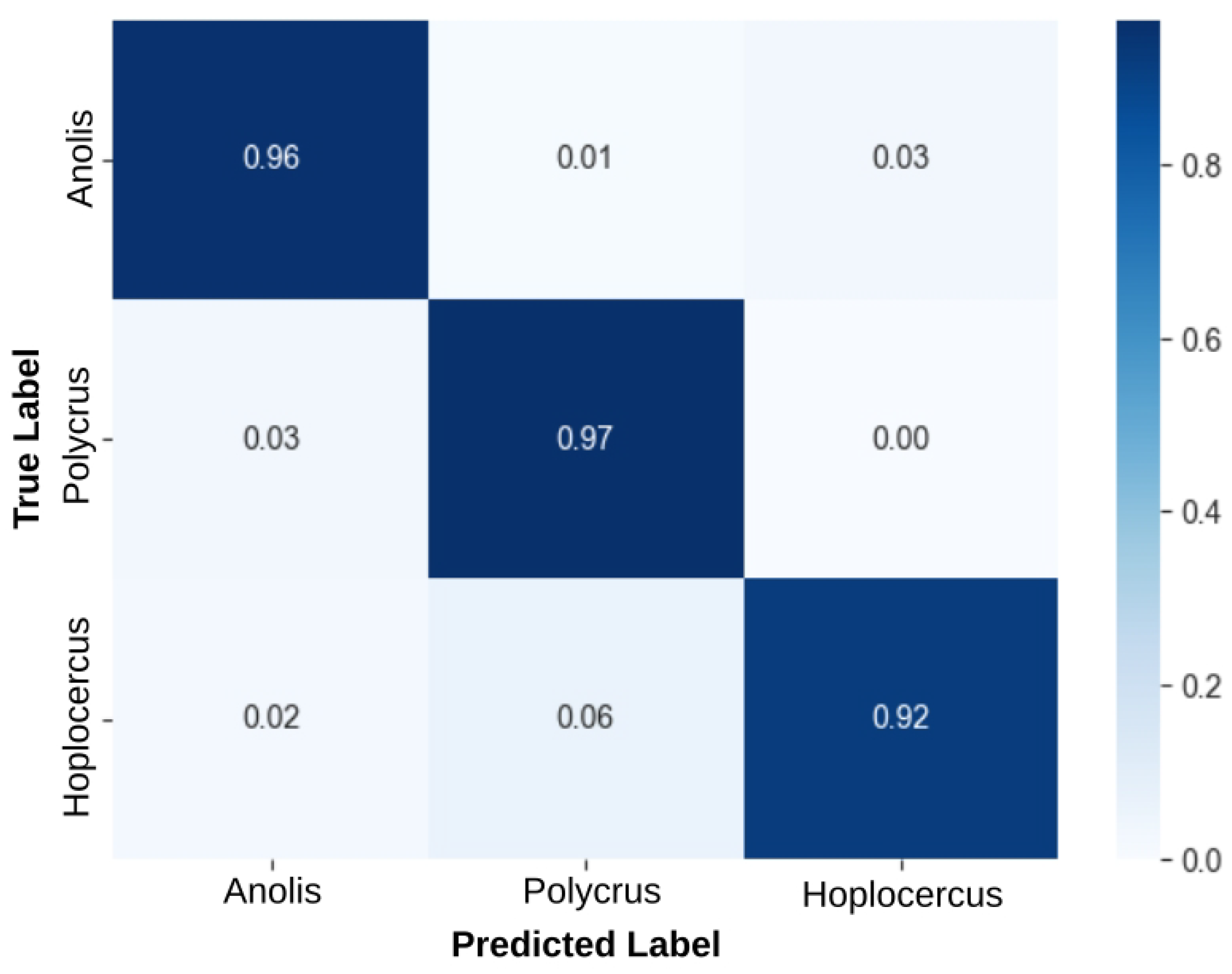
Model’s normalized confusion matrix. The confusion matrix for the best performing MobileNetV3-Small + Linear SVM model trained on the 20 top-ranked features dataset.

The performance in species classification by class proved to be highly efficient, as illustrated in Figure 5. Consequently, this ensures reliability in both accuracy and f1-score metrics. Furthermore, it is worth noting that there was little to no difference between these two metrics for the best model.

## Materials and methods

### Collection of 3D data samples

Data was collected at MPEG, located in Belém, Para, Brazil. MPEG is the second-oldest scientific research institution in Brazil, founded in 1866, and it houses a local herpetological collection with approximately 100,000 specimens of amphibians and reptiles [17]. Three species were selected for collection, namely: a) *Anolis fuscoauratus*; b) *Hoplocercus spinosus*; and c) *Polychrus marmoratus*; all species found in the Amazon region [18–20]. Figure 3 below shows pictures of individuals from each species.

**Fig 3.**
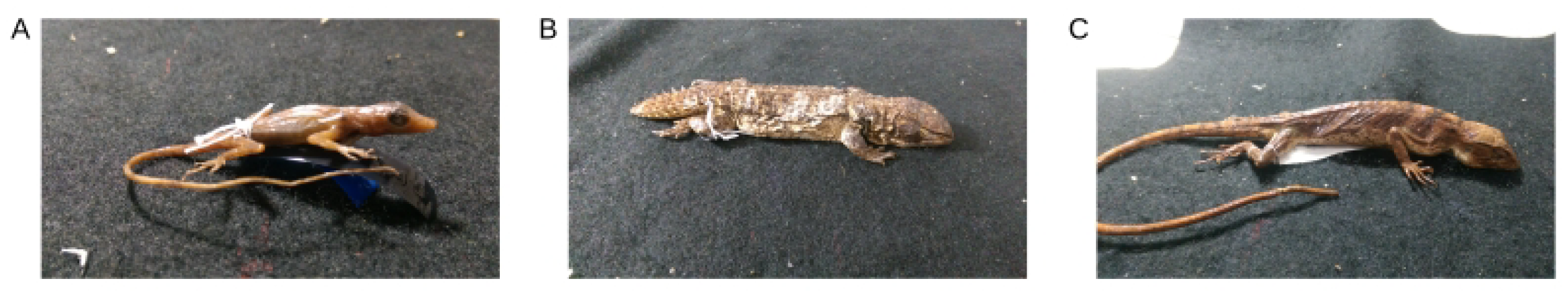
The three species selected for this study. (a) *Anolis fuscoauratus* (b) *Hoplocercus spinosus* (c) *Polychurs marmoratus*.

All specimens were preserved in alcohol, and the preservation conditions of each sample were a determining factor in selecting both the individuals and species chosen for this study. The selected individuals were then placed on a black cloth, and positioned on the collection bench to mitigate any visual noise that could interfere with identification. This simple strategy can be easily replicated in any environment, as in field data collection routines.

In recent studies using three-dimensional samples for species classification, the extensive use of Light Detection and Ranging (LiDAR), and Spectral Imaging (SI) are commonly used, particularly in studies using plants as specimens [21–23]. However, these technologies are costly and require highly specialized expertise, making them impractical for everyday use by experts in both laboratory and field settings. Furthermore, using not practical solutions such as LiDAR and SI makes it almost impossible to safely and easily reproduce the results, especially in areas where research funding is unstable.

As a solution, we adopted smartphone-based image capture from the dorsal, lateral, and ventral points of view to compose our samples. The use of smartphones offers a cost-effective alternative, enabling broader accessibility and usability for species classification. As can be seen in figure 4, three photos of each individual were taken, where each set of three images constitutes a single sample.

**Fig 4.**
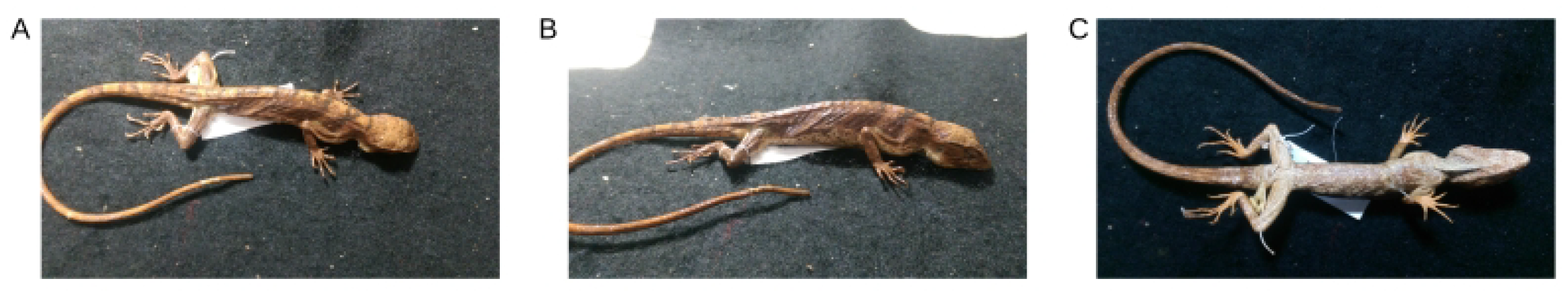
A sample comprised of the three points of view. A (a) dorsal, (b) lateral, and (c) ventral view of a *Polychurs marmoratus*, comprising one sample.

**Fig 5.**
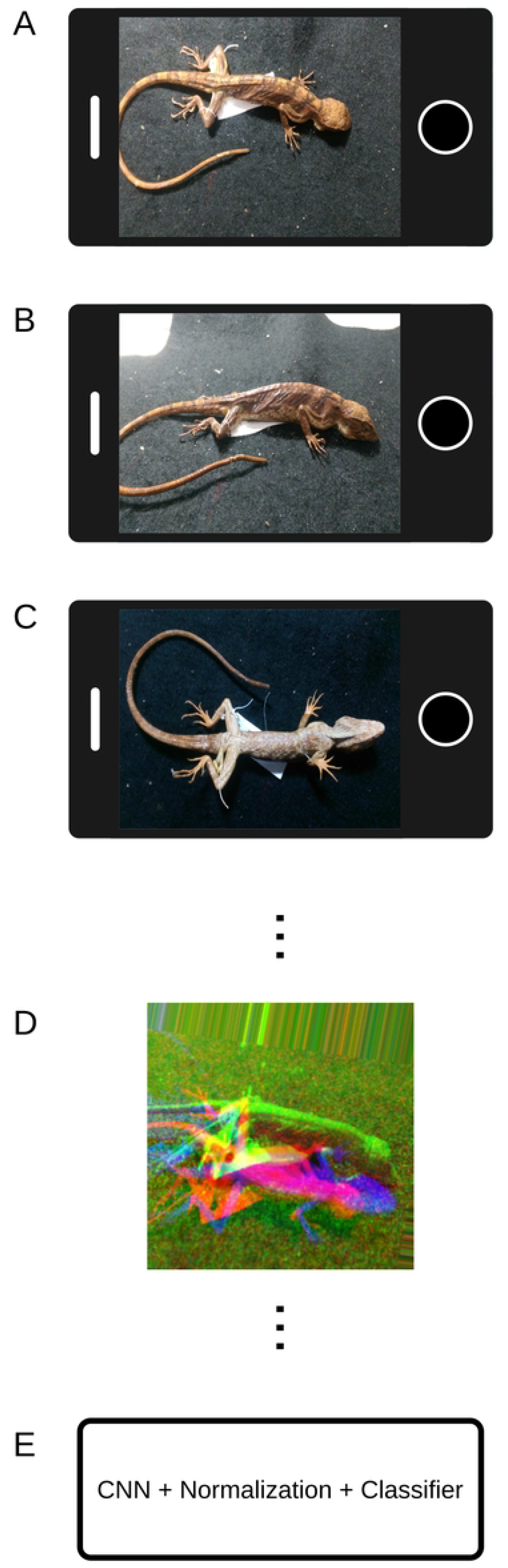
Classification pipeline for 3D representation of amazonian lizards. (a) take a photo of dorsal view (b) take a photo of lateral view (c) take a photo of ventral view (d) the three images are put together into one 3D sample (e) the model infers to what class that lizard belongs to.

It was necessary to remove images due to poor quality, a total of 80 three-dimensional samples, totaling 240 unique images, remained. Among these, there were 49 samples of *Anolis fuscoauratus*, 22 samples of *Hoplocercus spinosus*, and 9 samples of *Polychrus marmoratus*.

### Data samples processing

The first processing step was the organization of the samples with one image per RGB color channel, where dorsal = R, lateral = G, and ventral = B. Subsequently, all samples were resized to dimensions of 224 × 224 and standardized for the input layer of our CNN. The dataset was then divided into training and validation sets following an 80%-20% division, respectively, ignoring an additional hold-out validation set in favor of using cross-validation. We used TensorFlow’s (TF) image data generator module [24] for data augmentation, where random modifications such as Flip, Crop, Translate, etc., were applied to the samples without altering their fundamental characteristics, thus generating new synthetic samples in our dataset [25]. The outcome of data augmentation resulted in an increase from 80 initial three-dimensional samples to 3900 and 1790 in the training and testing sets, respectively.

### Deep learning models selection

We selected the class of MobileNet models for developing our species identification system. This class consists of highly efficient algorithms for mobile CV applications and embedded systems [26]. There are three main MobileNet models: a) MobileNet; b) MobileNetv2; and c) MobileNetV3, with the latter having two variants, namely: Large and Small [26–28].

The first model (MobileNet) is based on depth wise separable convolutions, which are a form of factorized convolutions that transform a regular convolution operation into depth wise, which significantly reduces both computational cost and model size [26]. The second model (MobileNetV2) introduces the new *inverted residual with a linear bottleneck* module [27], which expands to a higher dimension a compressed low-dimensional representation of the input data and then filters it using a lightweight depth wise convolution, reducing the memory requirements of the model. The third model (MobileNetV3) features an efficient redesign of the network architecture, coupled with a segmentation decoder that optimizes resource consumption for both of its variants, the Large, for devices with greater availability of resources, and the Small, for scenarios with more limited processing power [28].

We used and compared the performance of all available MobileNet network variants as feature extractors only. We did not retrain the models, and we appended a Global Average Pooling 2D layer at the end of each model for dimensionality reduction, and then we replaced their classification layers with classical ML algorithms. We adopted this hybrid approach because, there is evidence that using pre-trained models, such as MobileNets as feature extractors, can transfer their high accuracies acquired on ILSVRC to the new models they compose, without the need for computationally expensive retraining [14, 29, 30]. Moreover, the composition of a hybrid model with a classical algorithm serving as the final classifier drastically reduces the likelihood of the model presents overfitting [30].

### Machine learning models selection

The selection of classical ML algorithms was based on the criteria that it has to be commonly applied in research with biological databases [31], and pre-implemented in Scikit-learn (SKL) [32]. The chosen models were a) Support Vector Machine (SVM) with linear, rbf, poly kernels; b) K-Nearest Neighbors (KNN); c) Random Forest (RF); d) GaussianNB (GNB); and e) AdaBoost (ADB).

### Feature extraction process

From the original dataset we generated four new datasets of features, each one extracted with a different variation of MobileNet (V1, V2, V3-Large, and V3-Small), we call these full-features datasets.

### High-dimensional feature visualization

We applied the t-SNE on the full-features datasets. The t-SNE is a method which compresses a high-dimensional data on a two- or three-dimensional map [33], allowing us to understand the complexity of high-dimensional data visually.

### Dimensionality reduction process

We used the RF algorithm to compute a feature importance rank for each full-features dataset [34], then we used an importance score of 0.01 as a threshold to select the top-ranked features only, which amounted 20 columns for each full-features dataset. Now we have a 20 top-ranked features dataset for each corresponding full-features dataset.

### Machine learning models training & evaluation

For comparison, we first trained our ML models on each full-features dataset, and then on each 20 top-ranked features dataset. All datasets were normalized with *MinMaxScaler* [35]. It was a cross-validated training, with the *k-fold* and *random state* parameters set to 4, and 42, respectively. The models’ Accuracy and F1-Score were used for performance evaluation. We also used the McNemar’s statistical test for paired statistical model performance comparison [39].

### Best machine learning model optimization

We used a Bayesian Optimization (BO) process to improve the best ML model’s hyperparameters even further. By using BO, a surrogate for the model’s objective function is created, and a Gaussian Regressor quantifies the uncertainty for the surrogate [36]. The formula below shows the acquisition function Expected Improvement (EI), adopted in this study.

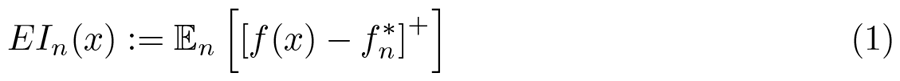

EI is popular due to its multi-modal nature and effective balance between exploration and exploitation of the search space for the best set of hyperparameters that will produce the lowest error on the model [37].

### Pipeline development & assembling

Our entire pipeline is open-source, and was developed using Python [38], the TF DL framework [24], and the SKL ML framework [32]. For image acquisition, we employed a replaceable smartphone HTC One M8, 32GB, Quad-Core 2.3GHz, with a 4MP× 2688×1520 440 ppi camera. The inference pipeline consists of five main stages: a) capturing a dorsal photo of the specimen; b) capturing a lateral photo of the specimen; c) capturing a ventral photo of the specimen; d) composing a three-dimensional sample from the aforementioned images; e) classifying the lizard species. Figure 5 below visually represents the pipeline sequence.

## Acknowledgments

We thank the Herpetology Laboratory from the Museu Paraense Emilio Goeldi for permitting us to collect the data used in this study. Our sincere appreciation also goes to the Instituto Tecnologico Vale for their support, providing full research scholarships for two of the authors, AGS and COB.

## References

1. Pyron, R Alexander and Burbrink Frank T and Wiens John J. A phylogeny and revised classification of Squamata, including 4161 species of lizards and snakes BMC evolutionary biology. 2013;13(1):1–54.

2. Stewart, Glenn R and Daniel Ronald S. Microornamentation of lizard scales: some variations and taxonomic correlations Herpetologica. 1975;117–130.

3. Villa, Alexander Gomez and Salazar, Augusto and Vargas, Francisco. Towards automatic wild animal monitoring: Identification of animal species in camera-trap images using very deep convolutional neural networks Ecological informatics. 2017;41:24–32.

4. Bell, Trent P and others. A novel technique for monitoring highly cryptic lizard species in forests Herpetological Conservation and Biology. 2009;4(3):415–425.

5. Miao, Zhongqi and Gaynor Kaitlyn M and Wang, Jiayun and Liu, Ziwei and Muellerklein, Oliver and Norouzzadeh Mohammad Sadegh and McInturff, Alex and Bowie Rauri CK and Nathan, Ran and Yu Stella X and others. Insights and approaches using deep learning to classify wildlife Nature Publishing Group UK London. 2019;9(1):8137.

6. Chen, R., Little, R., Mihaylova, L., Delahay, R., & Cox, R. Wildlife surveillance using deep learning methods Ecology and evolution. 2019;9(17):9453–9466.

7. Martineau, Maxime and Conte, Donatello and Raveaux, Romain and Arnault, Ingrid and Munier, Damien and Venturini, Gilles A survey on image-based insect classification Pattern Recognition. 2017;65:273–284.

8. Weinstein Ben G A computer vision for animal ecology Journal of Animal Ecology. 2018;87(3):533–545.

9. Waldchen, J and Mader, P. Machine learning for image based species identification Methods in Ecology and Evolution. 2018;9(11):2216–2225.

10. Bhardwaj, Manish and Singh, Anamika and Kumar, Manish and Sharma Sumit Kumar and Kumar, Jaideep Design and Analysis of Artificial Intelligence Model for the Global Issue of Poisonous Reptile Identification BioGecko. 2023;12(4):16–25.

11. Durso, Andrew M and Moorthy Gokula Krishnan and Mohanty Sharada P and Bolon, Isabelle and Salathe, Marcel and Ruiz de Castaneda, Rafael Supervised learning computer vision benchmark for snake species identification from photographs: Implications for herpetology and global health Frontiers in artificial intelligence. 2021;(4).

12. Binta Islam, Sazida and Valles, Damian and Hibbitts Toby J and Ryberg Wade A and Walkup Danielle K and Forstner Michael RJ Animal Species Recognition with Deep Convolutional Neural Networks from Ecological Camera Trap Images Multidisciplinary Digital Publishing Institute - Animals. 2023;13(9):1526.

13. Sharmin, Israt and Islam Nuzhat Farzana and Jahan, Israt and Ahmed Joye, Tasnem and Rahman Md Riazur and Habib Md Tarek Machine vision based local fish recognition SN Applied Sciences. 2019;(1):1–12.

14. Kornblith, Simon and Shlens, Jonathon and Le Quoc V Do better imagenet models transfer better? Proceedings of the IEEE/CVF conference on computer vision and pattern recognition. 2019;2661–2671.

15. Tuia, Devis and Kellenberger, Benjamin and Beery, Sara and Costelloe Blair R and Zuffi, Silvia and Risse, Benjamin and Mathis, Alexander and Mathis Mackenzie W and van Langevelde, Frank and Burghardt, Tilo and others Perspectives in machine learning for wildlife conservation Nature communications. 2022;13(1):792.

16. Norouzzadeh, Mohammad Sadegh and Morris, Dan and Beery, Sara and Joshi, Neel and Jojic, Nebojsa and Clune, Jeff A deep active learning system for species identification and counting in camera trap images Methods in ecology and evolution. 2021;12(1):150–161.

17. da Costa Prudente, Ana Lucia and da Cruz Ramos, Lorran Alves and da Silva Timoteo Monteiro and de Melo Sarmento, Joao Fabricio and Dourado Angelo Cortez Moreira and Silva Fernanda Magalhaes and de Almeida Paula Carolina Rodrigues and Dos Santos, Cleverson Rannieri Meira and de Sousa, Marcos Paulo Alves Dataset from the Snakes (Serpentes, Reptiles) collection of the Museu Paraense Emilio Goeldi, Para, Brazil Biodiversity Data Journal. 2019;7.

18. Vitt, Laurie J and Avila-Pires, Teresa Cristina S and Zani Peter A and Sartorius Shawn S and Esposito, Maria Cristina Life above ground: ecology of Anolis fuscoauratus in the Amazon rain forest, and comparisons with its nearest relatives Canadian Journal of Zoology. 2003;81(1):142–156.

19. Torres-Carvajal, Omar and de Queiroz, Kevin Phylogeny of hoplocercine lizards (Squamata: Iguania) with estimates of relative divergence times Molecular Phylogenetics and Evolution. 2009;50(1):31–43.

20. Murphy, John C and Lehtinen Rick M and Charles Stevland P and Wasserman, Danielle and Anton, Tom and Brennan, Patrick J Cryptic multicolored lizards in the Polychrus marmoratus Group (Squamata: Sauria: Polychrotidae) and the status of Leiolepis auduboni Hallowell Amphibian & Reptile Conservation. 2017.

21. Mayra, Janne and Keski-Saari, Sarita and Kivinen, Sonja and Tanhuanpaa, Topi and Hurskainen, Pekka and Kullberg, Peter and Poikolainen, Laura and Viinikka, Arto and Tuominen, Sakari and Kumpula, Timo and others Tree species classification from airborne hyperspectral and LiDAR data using 3D convolutional neural networks Remote Sensing of Environment. 2021;256:112–322.

22. Nezami, Somayeh and Khoramshahi, Ehsan and Nevalainen, Olli and Polonen, Ilkka and Honkavaara, Eija Tree species classification of drone hyperspectral and RGB imagery with deep learning convolutional neural networks Remote Sensing - MDPI. 2020;12(7):1070.

23. Polonen, Ilkka and Annala, Leevi and Rahkonen, Samuli and Nevalainen, Olli and Honkavaara, Eija and Tuominen, Sakari and Viljanen, Niko and Hakala, Teemu Tree species identification using 3D spectral data and 3D convolutional neural network 2018 9th Workshop on Hyperspectral Image and Signal Processing: Evolution in Remote Sensing (WHISPERS). 2018;1–5.

24. Martín Abadi, Ashish Agarwal, Paul Barham, Eugene Brevdo, Zhifeng Chen, Craig Citro, Greg S. Corrado, Andy Davis, Jeffrey Dean, Matthieu Devin, Sanjay Ghemawat, Ian Goodfellow, Andrew Harp, Geoffrey Irving, Michael Isard, Rafal Jozefowicz, Yangqing Jia, Lukasz Kaiser, Manjunath Kudlur, Josh Levenberg, Dan Mané, Mike Schuster, Rajat Monga, Sherry Moore, Derek Murray, Chris Olah, Jonathon Shlens, Benoit Steiner, Ilya Sutskever, Kunal Talwar, Paul Tucker, Vincent Vanhoucke, Vijay Vasudevan, Fernanda Viégas, Oriol Vinyals, Pete Warden, Martin Wattenberg, Martin Wicke, Yuan Yu, and Xiaoqiang Zheng TensorFlow: Large-Scale Machine Learning on Heterogeneous Systems Software available from tensorflow.org. 2015.

25. Xu, Mingle and Yoon, Sook and Fuentes, Alvaro and Park, Dong Sun A comprehensive survey of image augmentation techniques for deep learning Pattern Recognition. 2023;109347.

26. Howard, Andrew G and Zhu, Menglong and Chen, Bo and Kalenichenko, Dmitry and Wang, Weijun and Weyand, Tobias and Andreetto, Marco and Adam, Hartwig Mobilenets: Efficient convolutional neural networks for mobile vision applications arXiv preprint arXiv:1704.04861. 2017.

27. Sandler, Mark and Howard, Andrew and Zhu, Menglong and Zhmoginov, Andrey and Chen, Liang-Chieh Mobilenetv2: Inverted residuals and linear bottlenecks Proceedings of the IEEE conference on computer vision and pattern recognition. 2018;4510–4520.

28. Howard, Andrew and Sandler, Mark and Chu, Grace and Chen, Liang-Chieh and Chen, Bo and Tan, Mingxing and Wang, Weijun and Zhu, Yukun and Pang, Ruoming and Vasudevan, Vijay and others Searching for MobileNetV3 Proceedings of the IEEE/CVF international conference on computer vision. 2019;1314–1324.

29. Sowmya, M and Balasubramanian, M and Vaidehi, K Classification of Animals Using MobileNet with SVM Classifier Computational Methods and Data Engineering: Proceedings of ICCMDE. 2022;347–358.

30. Michele, Aurelia and Colin, Vincent and Santika Diaz D Mobilenet convolutional neural networks and support vector machines for palmprint recognition Procedia Computer Science. 2019;110–117.

31. Jovel, Juan and Greiner, Russell An introduction to machine learning approaches for biomedical research Frontiers in Medicine. 2021;8:771607.

32. Pedregosa, F. and Varoquaux, G. and Gramfort, A. and Michel, V. and Thirion, B. and Grisel, O. and Blondel, M. and Prettenhofer, P. and Weiss, R. and Dubourg, V. and Vanderplas, J. and Passos, A. and Cournapeau, D. and Brucher, M. and Perrot, M. and Duchesnay, E. Scikit-learn: Machine Learning in Python Journal of Machine Learning Research. 2011;12:2825–2830.

33. Van der Maaten, Laurens and Hinton, Geoffrey Visualizing data using t-SNE Journal of machine learning research. 2008;9(11).

34. Haq, Anwar Ul and Zhang, Defu and Peng, He and Rahman, Sami Ur Combining multiple feature-ranking techniques and clustering of variables for feature selection IEEE Access. 2019;17151482–151492.

35. Raju, VN Ganapathi and Lakshmi, K Prasanna and Jain Vinod Mahesh and Kalidindi, Archana and Padma, V Study the influence of normalization/transformation process on the accuracy of supervised classification Third International Conference on Smart Systems and Inventive Technology (ICSSIT). 2020;729–735.

36. Frazier, Peter I A tutorial on Bayesian optimization arXiv preprint arXiv:1807.02811. 2018.

37. Wang, Hao and van Stein, Bas and Emmerich, Michael and Back, Thomas A new acquisition function for Bayesian optimization based on the moment-generating function IEEE International Conference on Systems, Man, and Cybernetics (SMC). 2017:507–512.

38. Van Rossum, Guido and Drake Jr, Fred L Python Tutorial Centrum voor Wiskunde en Informatica Amsterdam, The Netherlands. 1995.

39. McCrum-Gardner, Evie Which is the correct statistical test to use? British Journal of Oral and Maxillofacial Surgery. 2008;46(1):38–41.

40. Pinho, Catarina and Kaliontzopoulou, Antigoni and Ferreira Carlos A and Gama, Joao. Identification of morphologically cryptic species with computer vision models: wall lizards (Squamata: Lacertidae: Podarcis) as a case study Zoological Journal of the Linnean Society. 2023;198(1):184–201.

41. Gray, Patrick C and Bierlich Kevin C and Mantell Sydney A and Friedlaender Ari S and Goldbogen Jeremy A and Johnston David W Drones and convolutional neural networks facilitate automated and accurate cetacean species identification and photogrammetry Methods in Ecology and Evolution. 2019;10(9):1490–1500

